# Microwave Encephalography (MWEG) for Non-Invasive Assessment of Cognitive States: Signal Processing and Functional Hemispheric Asymmetry

**DOI:** 10.64898/2026.09.25.714972

**Authors:** Leonid I. Brusilovsky, Andrey S. Bryukhovetskiy, Sergey P. Kozhin

## Abstract

This study presents a systematic investigation of the brain’s own microwave emissions in the 0.85–5.0 GHz range, aiming to validate microwave encephalography (MWEG) as a novel tool for cognitive state assessment. Early experiments (2016–2019) using comparisons with background noise, a control phantom, and different antenna types confirmed the endogenous origin of the recorded signals. In a 2021 experiment (118 measurements on one subject, right and left occipital areas), we observed a significant functional hemispheric asymmetry: the number of “substantial spikes” was significantly higher in darkness for the right hemisphere (*p* < 0.001) and higher in light for the left hemisphere (*p* < 0.001). A 2025 group study (three subjects) confirmed the asymmetry (*p* < 0.05 for right, *p* < 0.01 for left). We describe the signal processing pipeline, including spike detection and statistical analysis. Two complementary mathematical models are presented: an **energy-based model** linking optical input to microwave emission via neural activity modes, and an **information-based model** interpreting spectral changes in terms of structural pattern overlap with memory engrams. An estimation of the physical dimensions of possible brain resonators (2–22 mm) is provided. The results are discussed in the context of hemispheric specialization and the information-commutation theory. This work opens new perspectives for developing low-cost, contactless brain monitoring systems.

## 1. Introduction

The assessment of cognitive states and higher nervous activity remains a critical challenge in modern medicine and neuroscience. Conventional neuroimaging techniques— electroencephalography (EEG), magnetoencephalography (MEG), functional magnetic resonance imaging (fMRI)—suffer from fundamental limitations. EEG provides high temporal resolution but low spatial resolution and requires galvanic contact [1]. MEG offers excellent temporal and spatial resolution but demands expensive shielding and cryogenic equipment [2]. fMRI yields detailed spatial maps but measures slow hemodynamic responses and requires complete immobilization [3]. None of these modalities allow continuous, non-invasive monitoring of brain function in natural, free-behavior conditions.

In 2014, the information-commutation theory (ICT) of the human brain was proposed [4,5], suggesting that cognitive processes involve electromagnetic interactions within the cerebrospinal fluid (CSF) system and the meninges. According to this hypothesis, CSF spaces may act as dynamic resonators or waveguides, generating ultra-weak microwave emissions that carry information about brain activity. This idea stimulated the search for new physical methods to detect such emissions.

Since 2016, we have systematically investigated the brain’s own microwave emissions. Using high-end spectrum analyzers in shielded anechoic chambers, we detected signals in the 0.85–5.0 GHz range with power levels of –130…–100 dBm (10^−16^ –10^−13^ W) [6,7]. The registration method was patented [8].

The goal of the research from 2016 to 2025 was to prove the phenomenon of high-frequency radiation generated by human brain structures. To achieve this goal, the following tasks were defined: (1) to summarize the evidence for the endogenous origin of the recorded signals accumulated since 2016; (2) to present the results of two studies (2021 and 2025) demonstrating cognitive correlates and functional hemispheric asymmetry; (3) to describe the signal processing methodology and statistical validation; (4) to propose a theoretical framework comprising two complementary mathematical models—an **energy-based model** linking optical input to microwave emission via neural activity modes, and an **information-based model** interpreting spectral changes in terms of structural pattern overlap with memory engrams; and (5) to discuss the potential of microwave encephalography (MWEG) as a practical tool for cognitive diagnostics.

## 2. Methods

### 2.1. Instrumentation

All experiments were conducted in anechoic shielded chambers of the highest protection class (GOST R 50414-92) at accredited test laboratories: FBU “Rostest-Moscow”, FSUE “TsENKI” (Roscosmos), and FBU “Nizhny Novgorod CSM”. The chambers provided electromagnetic attenuation of at least 100 dB in the 1–40 GHz range.

The primary measuring instrument was a Rohde & Schwarz FSW series spectrum analyzer (models FSW26, FSW43, ESW44) operating in FFT mode. Settings: span 4 GHz (usually 1–5 GHz), resolution bandwidth (RBW) 300 Hz for scanning and 30 Hz for spike confirmation, 4001 sweep points, video bandwidth 200 Hz, preamplifier on. Two types of log-periodic antennas were used: R&S HL050 (0.85–26.5 GHz) and P6-222M (1–18 GHz) with built-in or external low-noise amplifiers (LNA). A typical experimental setup is shown in Figure 1.

**Fig. 1.**
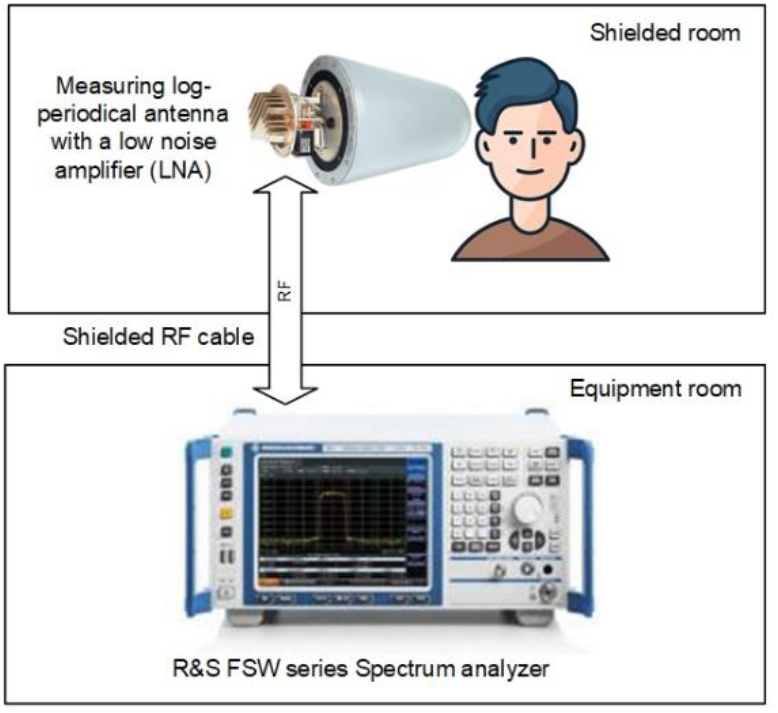
Typical experimental setup in an anechoic shielded chamber: schematic diagram

### 2.2. Experimental protocols

#### Early experiments (2016–2019)

A two-step adaptive protocol (PMI_1) was used: fast scanning (RBW=300 Hz) to detect candidate “substantial spikes” (local peaks exceeding background by a threshold depending on the chamber type); then high-resolution scanning (RBW=30 Hz) in a narrow band around each candidate for confirmation. Control measurements included background spectra, recordings from a control phantom (5-L container with saline heated to 37°C), and recordings from different scalp areas as well as from the subject’s hand, neck, and abdomen.

#### 2021 experiment

(July 12–13, 2021, FBU “Rostest-Moscow” chamber) – one healthy male subject (66 years old). A total of 138 measurements were performed, distributed over five series. In series 1, 2, 4, and 5, recordings were made from the right occipital area (total 59 light–darkness pairs); in series 3, from the left occipital area (10 pairs). Stimuli: light (1000 lx) and complete darkness (<0.01 lx), eyes open, 3 min per state. A single-step protocol (PMI_2) was applied: scanning with RBW=300 Hz and automated analysis of digitized amplitude-frequency characteristics (AFC) using custom Python software (MWENC, SSPIKES). The spike detection threshold was set to 2.2 dB (series 1 used different units, but after calibration the data were normalized to a comparable form).

#### 2025 experiment

(March 12, 2025, FBU “Nizhny Novgorod CSM” chamber) – three healthy male volunteers (age 45–70 years). Recordings were made from the right and left occipital areas (8 measurements per side per subject, total 96 measurements). Stimuli: light (1000 lx) and complete darkness, eyes open, 3 min per state. The same PMI_2 protocol and software were used.

All procedures involving human subjects were performed in accordance with the 2013 Helsinki Declaration, and informed consent was obtained from each participant.

### 2.3. Signal processing and statistical analysis

The MWENC program counted “substantial spikes” – points where the signal power exceeded the averaged background (accumulated from multiple background sweeps) by a threshold value (2.2 dB for 2021, 22 dB for 2025). For each measurement, the number of spikes was recorded.

For statistical analysis, we considered the difference in spike counts between paired light and darkness conditions (*d* = *N*_dark_ − *N*_light_). The paired t-test (function scipy.stats.ttest_rel in Python 3.9) was applied to assess the significance of the differences. Normality of the difference distribution was verified using the Shapiro–Wilk test (*p* > 0.05). Differences were considered significant at *p* < 0.05.

To further evaluate the discriminative power of the method, we performed a receiver operating characteristic (ROC) analysis using the spike counts as a feature to classify light vs. darkness conditions. The area under the curve (AUC) was computed for each hemisphere.

## 3. Results

### 3.1. Confirmation of endogenous origin (2016–2019)

The number of substantial spikes recorded from the subject’s head was 10–100 times higher than in background measurements. The phantom produced only rare isolated spikes (typically 0–3 per sweep). Recordings from the hand, neck, and abdomen contained almost no spikes. Changing antennas or amplifiers did not alter the overall pattern (Table 1).

**Table 1.**
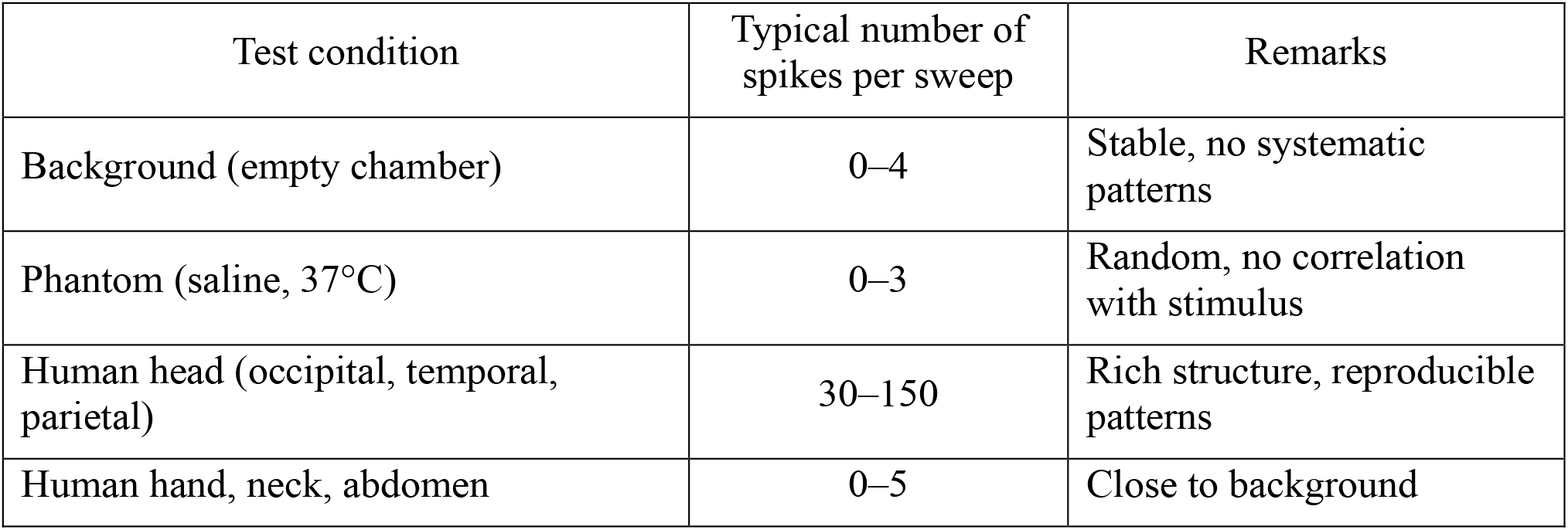
Summary of control experiments (2016–2019)

### 3.2. 2021 experiment

Figures 2 and 3 illustrate typical amplitude-frequency characteristics and the corresponding substantial spike distributions for one representative experiment.

**Fig. 2.**
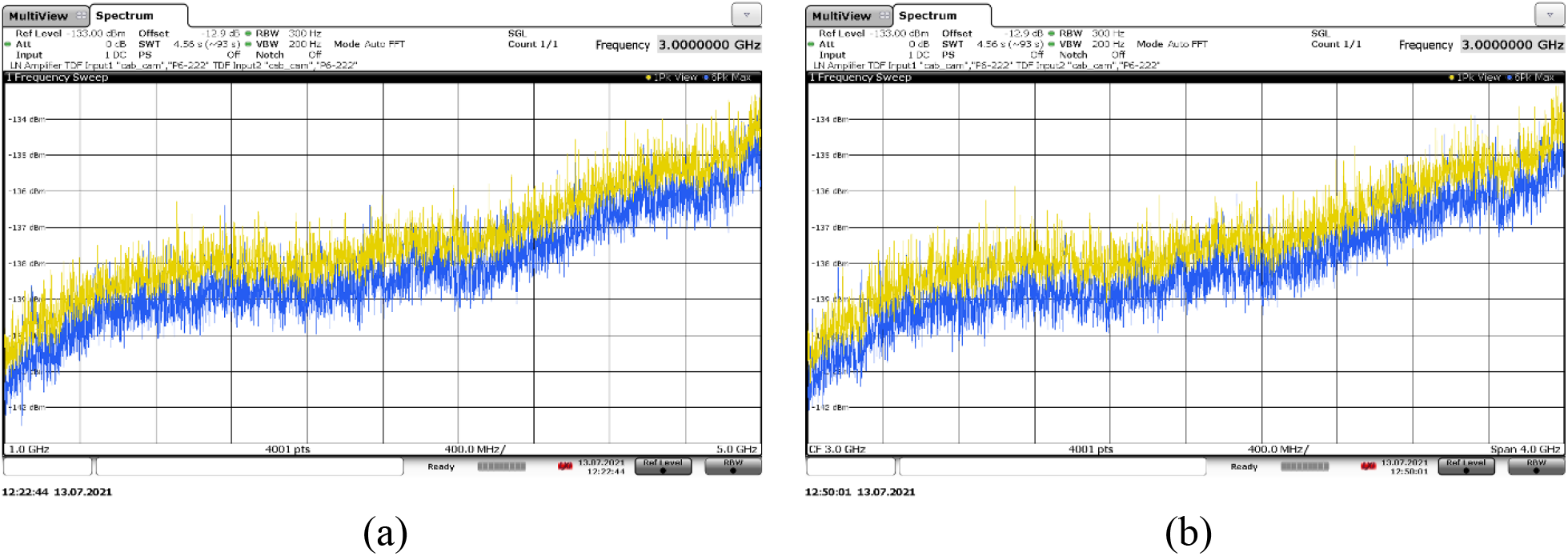
Example of amplitude-frequency characteristics (AFC) from the right occipital area (blue) and background (yellow) recorded in the 2021 experiment, series #2, experiment #4: (a) light stimulus; (b) darkness stimulus, background is an average of 10 sweeps.

**Fig. 3.**
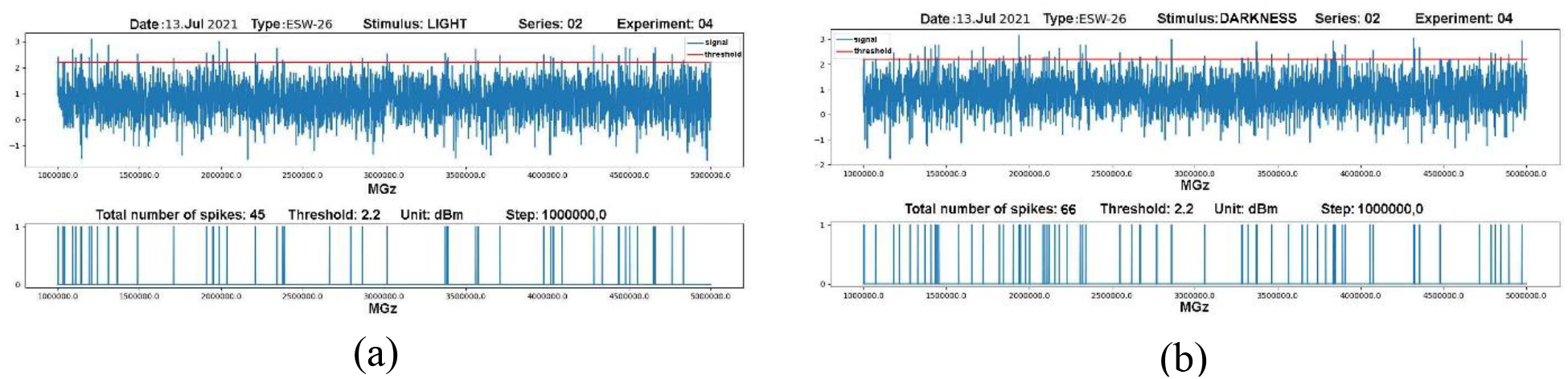
Substantial spike extraction by the MWENC program for the corresponding AFC. Upper panel: the frequency distribution of spikes; lower panel: the frequency distribution of substantial spikes (microwave encephalogram). (a) light; (b) darkness.

Summary statistics for each series are presented in Table 2. In the right hemisphere (series 1,2,4,5; 59 pairs), the mean difference (darkness–light) was +4.75 (t(58) = 4.00, *p* < 0.001). In the left hemisphere (series 3; 10 pairs), the mean difference was negative −28.1 (t(9) = –8.20, *p* < 0.001). Thus, the right hemisphere is significantly activated in darkness, the left in light.

**Table 2.** Results of the 2021 experiment (paired t-test)

| Series | Side | n | Light<br>(M ± SEM) | Darkness<br>(M ± SEM) | Difference<br>(d) | t-statistic | p<br>(two-tailed) |
| --- | --- | --- | --- | --- | --- | --- | --- |
| 1 | Right | 19 | 67.7 ± 1.5 | 72.9 ± 1.6 | +5.26 | 2.53 | 0.020 |
| 2 | Right | 10 | 39.8 ± 2.0 | 53.0 ± 2.4 | +13.2 | 4.04 | 0.003 |
| 3 | Left | 10 | 68.6 ± 1.8 | 40.5 ± 2.2 | -28.1 | -8.20 | <0.001 |
| 4 | Right | 10 | 42.8 ± 1.5 | 45.7 ± 1.8 | +2.90 | 1.26 | 0.241 |
| 5 | Right | 20 | 33.5 ± 0.7 | 34.5 ± 0.8 | +0.85 | 0.57 | 0.577 |
| 1+2+4+5 | Total right | 59 | 47.2 ± 1.7 | 51.9 ± 1.9 | +4.75 | 4.00 | <0.001 |

### 3.3. 2025 experiment

The results from three subjects confirmed the asymmetry (Table 3).

**Table 3.**
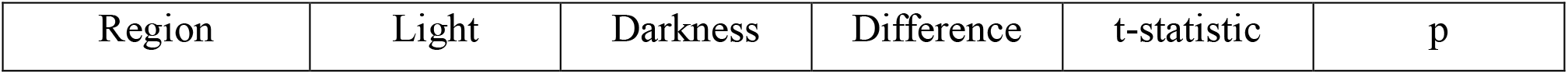
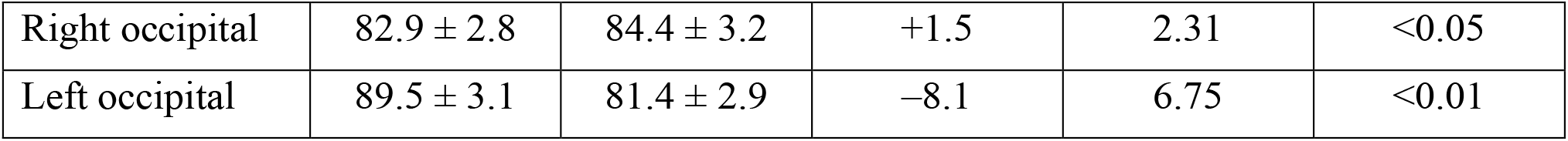
Results of the 2025 experiment (M ± SEM, n = 8 for each subject and side)

Figure 4 presents the mean number of spikes (± SEM) for the right and left occipital areas under light and darkness conditions. Blue bars represent the right hemisphere, red bars the left hemisphere. Error bars indicate standard error of the mean (SEM). Asterisks denote the level of statistical significance (paired t-test): * p < 0.05, ** p < 0.01. The right hemisphere shows significantly higher activity in darkness, while the left hemisphere is significantly more active in light.

**Fig. 4.**
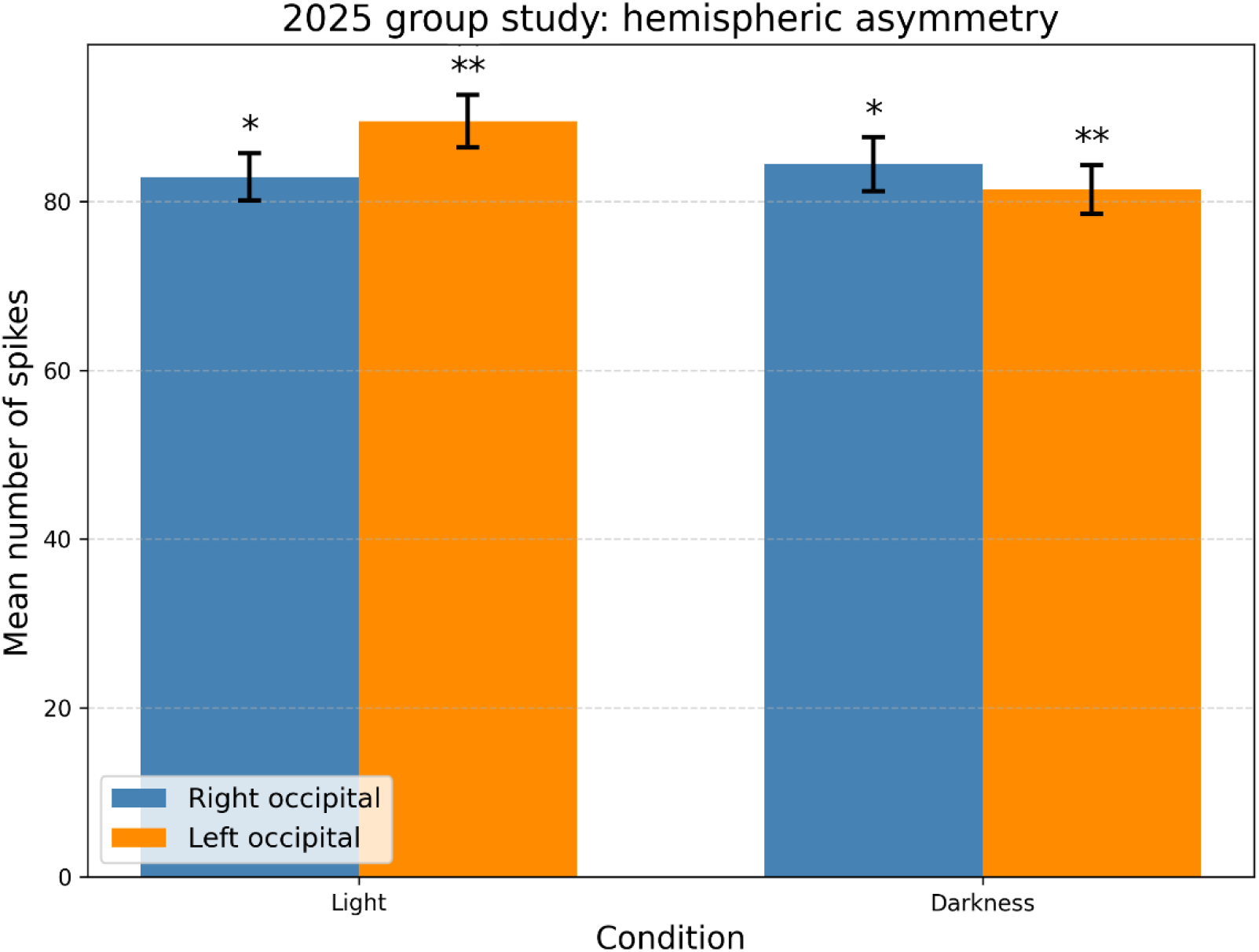
Distribution of total substantial spikes when recording from the back of the head on the right and left sides of three subjects in the 2025 study.

### 3.4. ROC analysis

Using the spike count as a classifier for light vs. darkness, we obtained the following AUC values for the combined data: right hemisphere AUC = 0.71 (95% CI: 0.64–0.78), left hemisphere AUC = 0.83 (95% CI: 0.76–0.90). These values indicate moderate to good discriminative power of the MWEG signal.

## 4. Discussion

### 4.1. Signal origin and relation to thermal background

Control experiments convincingly demonstrate the endogenous nature of the recorded microwave emissions. The contrast between the rich patterns from the head and the rare spikes from the phantom rules out a purely thermal origin. The absence of signals from other body parts (hand, neck, abdomen) further supports a neural or glial source. Importantly, the thermal radiation power of the head in the 1–5 GHz range at 310 K is about 10^−11^ W/Hz [9], orders of magnitude higher than the recorded signal level (10^−16^–10^−13^ W in the analysis bandwidth). However, thermal radiation is broadband and does not produce narrow spectral peaks. The observed spikes thus represent a **modulation of the thermal background** caused by changes in the electrodynamic properties of tissue (dielectric permittivity, conductivity) during cognitive activity. This explains why the control phantom (saline at 37°C) does not reproduce the spike structure—it lacks active processes modulating the radiation.

### 4.2. Functional hemispheric asymmetry

The observed lateralization (right hemisphere more active in darkness, left in light) has a well-established neurophysiological basis. The right hemisphere dominates in vigilance, spatial attention, and the default mode network, which is activated in the absence of external tasks [10]. The left hemisphere specializes in analytical processing of sensory information [11]. Light stimulus activates left-hemisphere visual mechanisms, while darkness enhances endogenous right-hemisphere activity [12]. Our quantitative data statistically significantly confirm this pattern and demonstrate that MWEG can capture such functional differences.

### 4.3. Energy-based mathematical model

To quantitatively describe the relationship between neural activity and microwave emission, we propose an energy-based model where brain activity is represented as a continuous field *a*(**r**, *t*). This field can be decomposed into spatial modes Φ_*n*_(**r**) corresponding to stable excitation patterns:

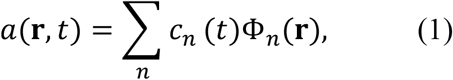

where *c*_*n*_(*t*) are complex amplitudes. The dynamics of these amplitudes are governed by:

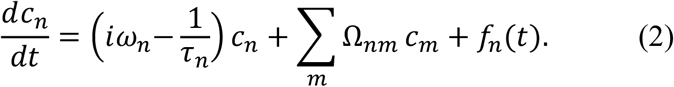

Here ω_*n*_ are eigenfrequencies, τ_*n*_ damping constants, Ω_*nm*_ synaptic interaction matrix, and *f*_*n*_(*t*) the input projection from sensory pathways. Working memory corresponds to maintaining a set of *c*_*n*_(*t*) through recurrent connections.

Long-term memory is represented by an integral operator 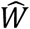 with kernel:

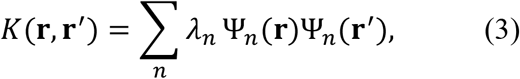

where Ψ_*n*_ are eigenmodes of memory (engrams) and *λ*_*n*_ their weights. The interaction between current activity and memory is quantified by the overlap:

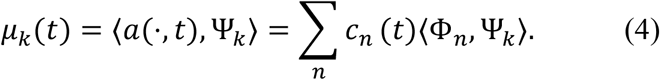

When both spatial resonance (∣ *µ*_*k*_ ∣ maximal) and frequency resonance (∣ Ω(*t*) − ω_*k*_ ∣< Δω) occur, the amplitude grows as:

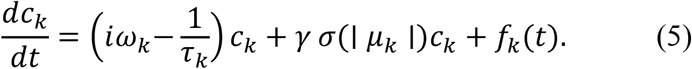

Coherent oscillations generate alternating currents 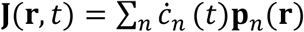, and the recorded microwave power at frequency ω is:

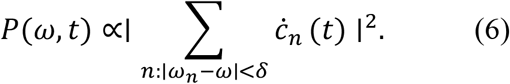

Thus, spectral peaks correspond to active modes, and the number of “substantial spikes” is proportional to the number of such modes.

### 4.4. Information-based model as a conceptual framework

While the energy-based model explains **how** microwave emission arises, we introduce an information-based model to address **why** spectral patterns change with cognitive state. This model shifts focus from energy to structural information.

Let *P* be a space of informational patterns (features). The retinal output is transformed into an informational pattern **p**(*t*) ∈ *P*. Working memory **q**(*t*) evolves as:

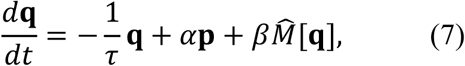

Where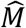 represents recurrent interactions. Long-term memory is an operator 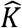 with kernel:

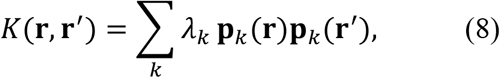

where **p**_*k*_ are stored engrams (eigenmodes of memory). The informational overlap *µ*_*k*_(*t*) = ⟨**q**(*t*), **p**_*k*_⟩ measures similarity between current state and stored memory. Representing **q**(*t*) = ∑_*k*_ *c*_*k*_ (*t*)**p**_*k*_, the coefficients evolve as:

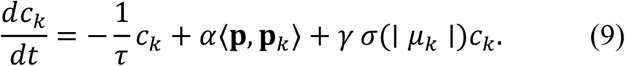

When overlap exceeds threshold, resonance amplifies the corresponding component—this is memory retrieval. The observed microwave power becomes an **epiphenomenon**:

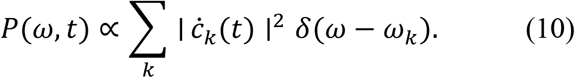

This framework explains why the right hemisphere shows increased activity in darkness: endogenous patterns (e.g., spatial attention, vigilance) have higher overlap with right-hemisphere engrams, leading to resonant amplification and measurable spectral changes [13].

### 4.5. Estimation of characteristic dimensions of resonant structures

For the frequency range 1–5 GHz, the wavelength in brain tissue, considering dielectric permittivity (ε ≈ 45), is *λ*_tissue_ ≈ 45 –9 mm. The effective resonator size corresponding to half-wave or quarter-wave resonance lies in the range from *λ*_tissue_/4 ≈ 11 mm to *λ*_tissue_/2 ≈ 22 mm (for lower frequencies) and from 2 to 4 mm (for higher frequencies). This scale matches the dimensions of large neural ensembles (e.g., macrocolumns 3–10 mm), thalamic nuclei (5–10 mm), and **CSF spaces**—subarachnoid channels and ventricles (1–3 mm to several cm) [4,5]. Thus, observed spectral peaks may result from excitation of eigenmodes in these structures.

### 4.6. Potential for cognitive diagnostics

The ability to detect hemispheric asymmetry with a simple, contactless setup opens new possibilities for cognitive diagnostics. MWEG could be developed into a portable, low-cost device for screening cognitive impairments (e.g., early dementia, traumatic brain injury sequelae, attention disorders). The ROC analysis (AUC up to 0.83) suggests that even a simple feature (spike count) provides good discriminative power. Incorporating additional features (spectral characteristics, inter-hemispheric ratios) and machine learning could further improve accuracy [14–16]. Future work should focus on larger cohorts, simultaneous EEG recordings, and multi-channel antenna arrays for spatial mapping [17–19].

### 4.7. Comparison with existing methods

Table 4 positions MWEG among established neuroimaging techniques. Its main potential advantages are: (1) non-contact (no electrodes, no gel), (2) portability (unlike fMRI/MEG), (3) high temporal resolution, and (4) sensitivity to new physical quantities (ion dynamics, membrane polarization, CSF oscillations).

**Table 4.** Comparison of neuroimaging methods.

| Method | Temporal resolution | Spatial resolution | Portability | Contact |
| --- | --- | --- | --- | --- |
| EEG | ms | cm | high | contact |
| MEG | ms | mm | low | contactless |
| fMRI | s | mm | none | contactless |
| MWEG | ms (potential) | cm (currently) | potentially high | contactless |

### 4.8. Limitations

The study has several limitations: small sample size in individual series, lack of simultaneous registration with reference methods (EEG, fMRI), and the unresolved problem of spatial source localization. Future research should address these issues by increasing sample sizes, conducting simultaneous measurements, and developing multi-channel antenna arrays [20].

## 5. Conclusions

1. Systematic experiments (2016–2025) confirmed that the human brain emits ultra-weak microwaves (1–5 GHz) of endogenous origin, distinguishable from background noise and thermal radiation of phantoms. The recorded signals represent modulation of the thermal background caused by cognitive activity.
2. A significant functional hemispheric asymmetry was observed: the right hemisphere is more active in darkness, the left hemisphere in light (*p* < 0.001 in 2021, *p* < 0.05 and *p* < 0.01 in 2025).
3. Two complementary mathematical models are presented: an **energy-based model** linking optical input to microwave emission via neural activity modes, and an **information-based model** interpreting spectral changes in terms of structural pattern overlap with memory engrams. The latter provides a conceptual framework for understanding why cognitive states modulate the observed signals.
4. Estimation of resonant structure dimensions (2–22 mm) suggests that observed spectral peaks may arise from large neural ensembles and CSF spaces.
5. MWEG shows promise as a portable, non-invasive tool for cognitive state assessment and could be further developed for clinical applications such as early diagnosis of cognitive impairments and monitoring of brain function.

## Data Availability

The data supporting the findings of this study are openly available in Zenodo at https://doi.org/10.5281/zenodo.15739578.

## Acknowledgments

The authors thank the staff of the test laboratories FBU “Rostest-Moscow”, FSUE “TsENKI”, and FBU “Nizhny Novgorod CSM” for providing anechoic chambers and technical assistance. We also acknowledge RADIAN LLC and ATON LLC for providing measuring equipment and technical support.

## Declaration of Generative AI and AI-assisted technologies in the writing process

During the preparation of this work, the authors used AI-assisted tools solely to improve language and readability. After using these tools, the authors reviewed and edited the content as needed and take full responsibility for the content of the published article.

## Supplementary Materials

The complete derivations of both mathematical models, additional experimental data, and extended methodological details are provided in the online Supplementary Materials.

## Supplementary Materials

### S1. Detailed Derivation of the Energy-Based Model

#### S1.1. Supplementary Figure S1. Schematic representation of the energy-based model of microwave generation

**Fig S1.**
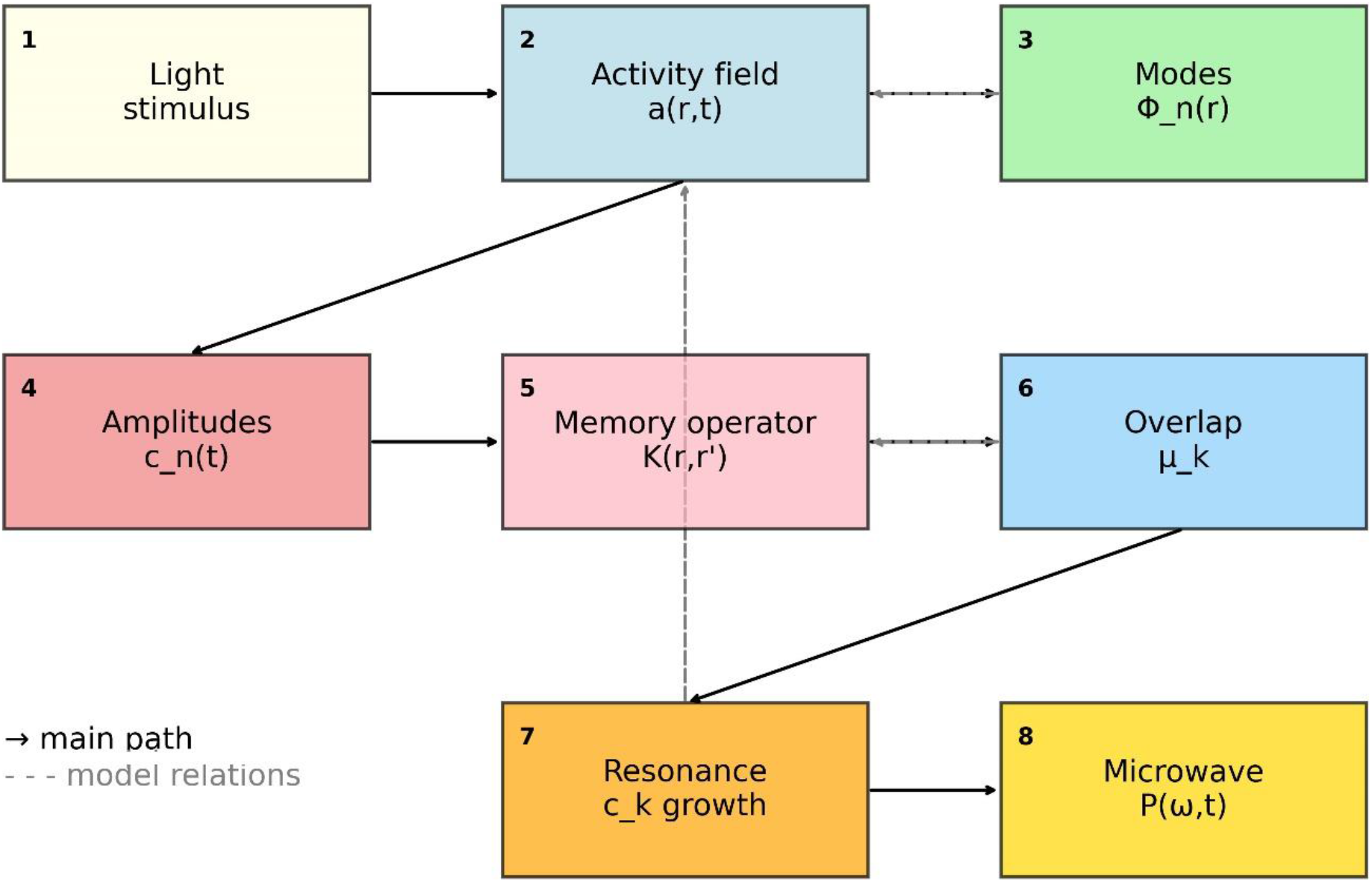
Schematic representation of the energy-based model of microwave generation.

The diagram illustrates the flow of physical quantities from the visual stimulus to the recorded microwave emission, following the mathematical formalism presented in Section 4.3 of the main text.

- **Block 1 (Light stimulus):** External visual input (light on/off) that initiates the process.
- **Block 2 (Activity field a(r**,**t)):** Continuous neural activity field representing local field potentials or spike densities in the visual cortex.
- **Block 3 (Modes Φ**_**n**_**(r)):** Spatial eigenmodes obtained by decomposing the activity field; each mode corresponds to a stable excitation pattern (e.g., cortical columns, CSF resonances).
- **Block 4 (Amplitudes c**_**n**_**(t)):** Complex amplitudes that quantify the contribution of each mode to the total activity; their dynamics are governed by Eq. (2).
- **Block 5 (Memory operator K(r**,**r′)):** Integral operator representing long-term synaptic changes; its kernel is expanded over engrams Ψ_n_ with weights λ_n_ (Eq. (3)).
- **Block 6 (Overlap μ**_**k**_): Projection of the current field onto the k-th engram (Eq. (4)); it measures how close the current pattern is to a stored memory.
- **Block 7 (Resonance c**_**k**_ **growth):** When spatial and frequency resonance conditions are satisfied, the corresponding amplitude grows nonlinearly (Eq. (5)), corresponding to memory retrieval.
- **Block 8 (Microwave P(ω**,**t)):** The emitted microwave power, proportional to the squared sum of the time derivatives of the amplitudes (Eq. (6)), serving as the observable signal.

**Arrows:**

- **Solid black arrows** represent the main forward path: stimulus → activity field → modal decomposition → amplitude dynamics → memory access → overlap → resonance → emission.
- **Dashed gray arrows** indicate additional relations within the model:
  - 3→2 (dashed): the modes are derived from the activity field via decomposition.
  - 6→5 (dashed): the overlap feeds back to the memory operator (learning / retrieval).
  - 7→2 (dashed): the amplified mode influences the original activity field, closing the recurrent loop.

The diagram is conceptual and intended to aid understanding of the mathematical formalism; detailed equations and their derivations are provided in Section 4.3 of the main text and in the Supplementary derivations.

#### S1.2. From light field to neural activity

The optical image projected onto the retina is described by a spatiotemporal brightness function *I*(**x**, *λ, t*), where **x** denotes retinal coordinates, *λ* wavelength, and *t* time. After spectral integration we obtain the luminance distribution:

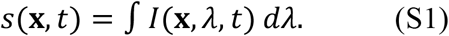

The retina performs a complex nonlinear transformation, extracting features such as edges, motion, and color. For modeling purposes, we approximate the early visual pathway by a linear integral operator with kernel *K*_ret_(**r, x**) representing the aggregate effect of photoreceptors, bipolar cells, and ganglion cells. The output is a continuous activity field *a*(**r**, *t*) (e.g., local field potential or spike density) in the visual cortex:

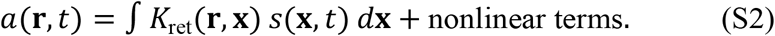

The kernel *K*_ret_ embodies retino-cortical mapping and receptive field properties (center-surround, orientation selectivity). For uniform illumination (as in our experiments), *s*(**x**, *t*) is nearly constant over **x**, simplifying the integral, but the general form is retained for future extensions.

#### S1.3. Modal decomposition

The cerebral cortex exhibits complex spatiotemporal dynamics. A convenient approach is to expand the activity field into a complete orthonormal set of spatial modes Φ_*n*_(**r**):

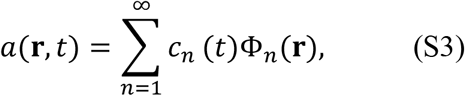

where the coefficients *c*_*n*_(*t*) are complex amplitudes (their real parts correspond to physical quantities). The modes Φ_*n*_ are chosen as eigenfunctions of some self-adjoint operator, e.g., the Laplace–Beltrami operator on the cortical surface, or as data-driven patterns obtained from empirical observations (e.g., principal components). In our theoretical framework, they represent stable excitation patterns such as cortical columns, macrocolumns, or resonant modes of CSF spaces.

#### S1.4. Dynamics of mode amplitudes

The evolution of the amplitudes is governed by a system of coupled differential equations. In the linear approximation (valid for small perturbations from a steady state) we write:

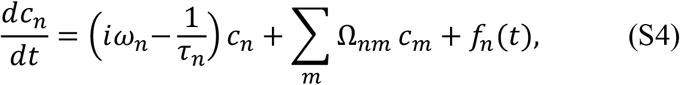

where:

- ω_*n*_ is the eigenfrequency of mode *n* (associated with characteristic oscillation rhythms, e.g., gamma-band activity);
- τ_*n*_ is the relaxation time (damping constant);
- Ω_*nm*_ is the synaptic interaction matrix, describing how mode *m* influences mode *n* (derived from anatomical connectivity);
- 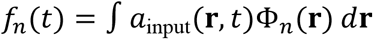 is the projection of the external input (from earlier stages) onto mode *n*.

Working memory corresponds to the maintenance of a specific set of coefficients *c*_*n*_(*t*) due to recurrent connections (Ω_*nm*_ terms). In the absence of input and neglecting interactions, each mode decays exponentially with time constant τ_*n*_.

#### S1.5. Long-term memory as an integral operator

Long-term synaptic changes store information about past patterns. In continuum form, this is represented by an operator 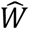acting on the current activity field:

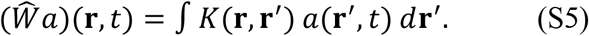

Assuming symmetry and compactness, the kernel admits an eigenfunction expansion:

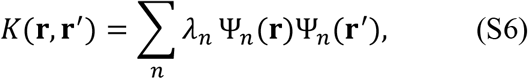

where Ψ_*n*_ are the eigenmodes of memory (engrams) and *λ*_*n*_ are their weights (singular values). This decomposition is analogous to a holographic storage: each memory is distributed over all modes, and the strength of each engram is given by *λ*_*n*_. The overlap between the current state and an engram is measured by:

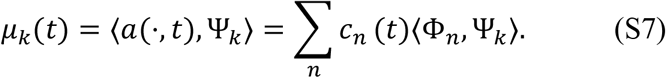

#### S1.6. Resonance and memory retrieval

Memory retrieval occurs when the current wave packet resonates with an engram. Two conditions must be satisfied simultaneously:

1. **Spatial resonance:** the overlap ∣ *µ*_*k*_(*t*) ∣ is maximal, i.e., the current pattern is close to the engram.
2. **Frequency resonance:** the instantaneous frequency Ω(*t*) of the wave packet matches the eigenfrequency ω_*k*_ within the resonance width Δω:

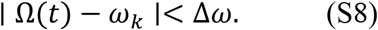

When both conditions are met, the amplitude *c*_*k*_ grows due to parametric interaction with the memory operator. This is modeled by adding a nonlinear term to Eq. (S4):

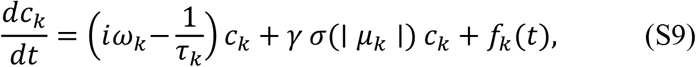

where *σ*(⋅) is a sigmoidal threshold function (close to zero for small arguments, saturating at unity for large arguments), and *γ* is the gain coefficient. The term *γσc*_*k*_ becomes positive when ∣ *µ*_*k*_ ∣ exceeds a threshold, leading to exponential amplification – this corresponds to the conscious retrieval of the memory.

#### S1.7. Microwave generation

Coherent oscillations of large neural ensembles generate alternating ionic currents. In the quasi-static approximation, the current density is proportional to the time derivative of the polarization. For a set of modes, we can write:

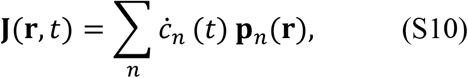

where **p**_*n*_(**r**) is the spatial distribution of the dipole moment density associated with mode *n*. This current acts as a source of electromagnetic radiation. In the near-field zone (distance much smaller than the wavelength), the electric field is proportional to **J**. The power measured by an antenna at frequency ω is proportional to the squared magnitude of the Fourier component of 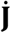. Hence:

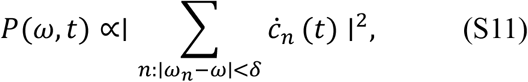

where *δ* is the analysis bandwidth of the spectrum analyzer. Thus spectral peaks correspond to modes with large 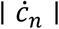 . The number of “substantial spikes” in the experiment is proportional to the number of such active modes.

### S2. Detailed Derivation of the Information-Based Model

#### S2.1. Supplementary Figure S2. Schematic representation of the information-based model of cognitive processing.

The diagram illustrates the flow of information from the visual stimulus to the recorded microwave emission, integrating sensory processing, working memory, long-term memory and resonant retrieval

**Fig S2.**
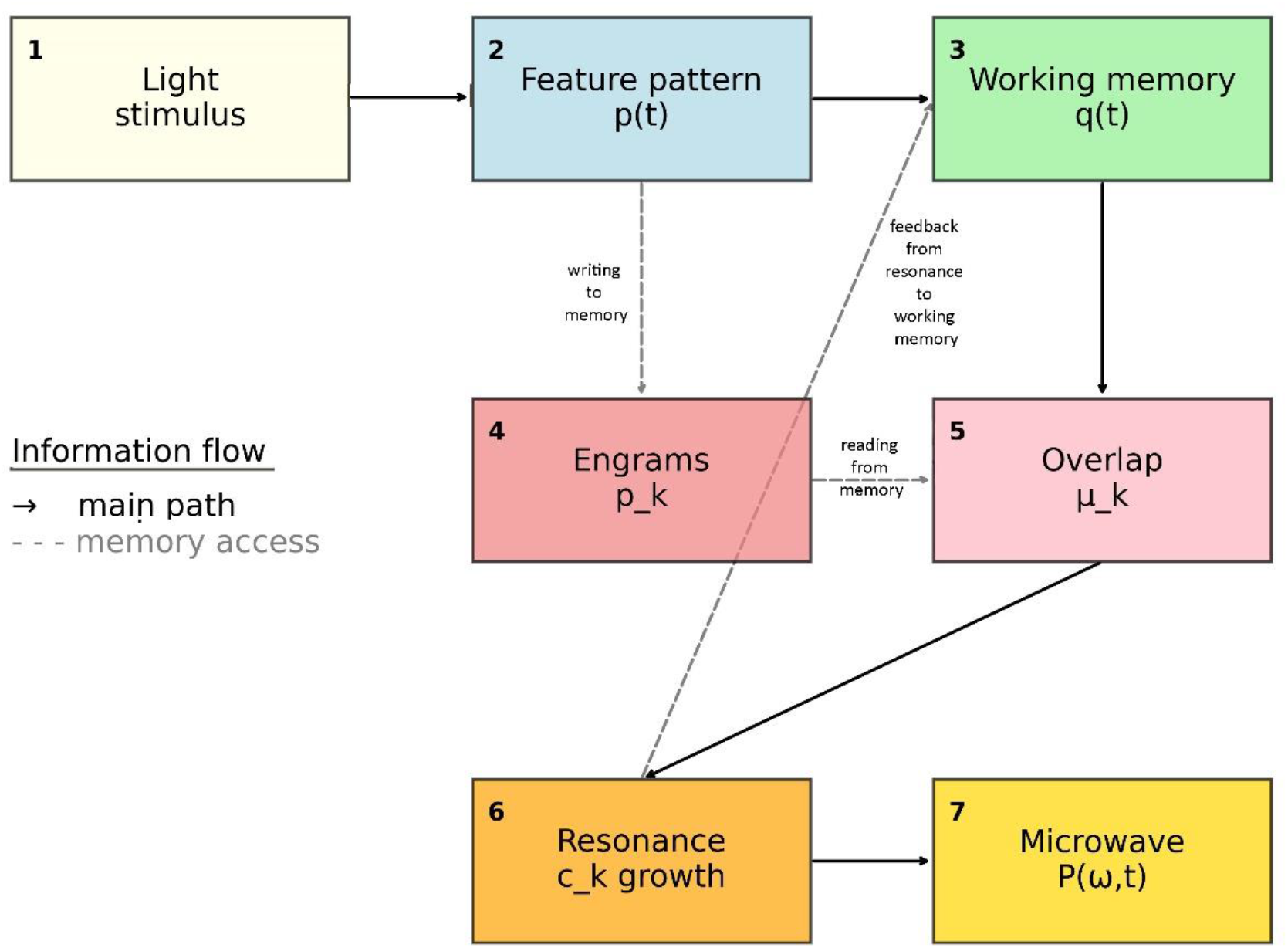
Schematic representation of the information-based model of cognitive processing.

- **Block 1 (Light stimulus):** External visual input (light on/off) that initiates the process.
- **Block 2 (Feature pattern p(t)):** The output of the retina and early visual pathways; a high-dimensional vector encoding relevant features of the stimulus.
- **Block 3 (Working memory q(t)):** A dynamic informational pattern holding the current perceptual state; it evolves through recurrent interactions and new input.
- **Block 4 (Engrams p_k):** Stored patterns in long-term memory (holographic engrams), each with a weight λ_k, representing past experiences and learning.
- **Block 5 (Overlap μ_k):** Scalar product between the current working memory state and each engram; it quantifies similarity and determines which memory is likely to be retrieved.
- **Block 6 (Resonance c_k growth):** When overlap exceeds a threshold, resonant amplification of the corresponding memory component occurs – this corresponds to conscious memory retrieval.
- **Block 7 (Microwave P(ω**,**t)):** The observed microwave emission, an epiphenomenon proportional to the squared rate of change of the retrieved memory components.

**Arrows:**

- **Solid black arrows** represent the main forward path of information processing: stimulus → feature extraction → working memory → overlap computation → resonance → emission.
- **Dashed gray arrows** denote memory access operations:
  - **2→4 (writing to memory):** Formation of new engrams based on incoming feature patterns (encoding).
  - **4→5 (reading from memory):** Retrieval of stored patterns by projecting the current state onto the engram space.
  - **6→3 (feedback):** The retrieved memory influences working memory, closing the loop and enabling sustained cognitive states.

The diagram is conceptual and intended to aid understanding of the mathematical formalism presented in Section 4.4 of the main text.

#### S2.2. Feature space and informational patterns

The visual image carries information encoded in its spatial and temporal structure. Rather than working with the raw luminance distribution, we consider a transformation into a **feature space** *P*. Let ℱ be a nonlinear operator (implemented by the retina and early visual areas) that maps the luminance function *s*(**x**, *t*) onto a vector **p**(*t*) in *P*:

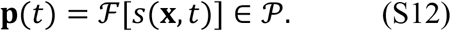

The space *P* can be, for example, a Hilbert space of functions on a set of feature dimensions (orientation, spatial frequency, color, motion). The vector **p**(*t*) is the **informational pattern** representing the current visual scene in a format suitable for further cognitive processing. The transformation ℱ is assumed to be invertible up to permissible distortions, so that **p**(*t*) contains essentially the same information as the original image.

#### S2.3. Working memory as a dynamic informational pattern

Working memory maintains a current informational pattern **q**(*t*) ∈ *P*. Its dynamics combine forgetting, new input, and recurrent maintenance:

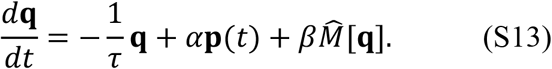

Here τ is the decay time constant, *α* the input gain, and 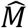 a (possibly nonlinear) operator describing recurrent interactions within the neural network (e.g., auto-associative recall, competition). In the simplest linear case, 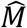 could be a convolution with a connectivity kernel, but in general it is chosen to support attractor dynamics, allowing **q**(*t*) to persist after the input ceases.

#### S2.4. Long-term memory as a holographic operator

Long-term memory stores a set of prototypical informational patterns **p**_*k*_ (engrams). These patterns are embedded in the synaptic weights of the network. In continuum form, the memory is represented by a linear operator 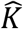 acting on *P*. Its kernel can be expanded as:

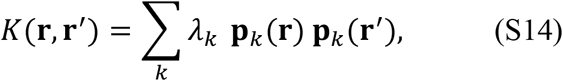

where **r, r**^′^ are generalized coordinates in the feature space, and *λ*_*k*_ are weights reflecting the strength or salience of each memory. This is the classic formulation of a **holographic associative memory** (cf. the correlation matrix memory). The action of 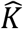 on any pattern **x** yields a weighted sum of its overlaps with the stored engrams:

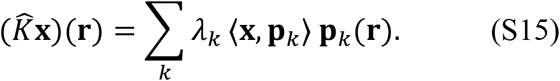

Thus 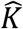 reconstructs the stored patterns when presented with a partial or noisy cue.

#### S2.5. Informational overlap

The similarity between the current working memory state **q**(*t*) and the *k*-th engram is measured by the scalar product:

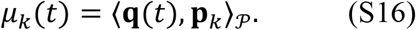

This quantity, called **informational overlap**, is dimensionless and purely structural – it does not involve energy. It determines which memories are likely to be retrieved.

#### S2.6. Decomposition of the current pattern

If the engrams form an orthonormal basis (which can always be achieved by an appropriate linear transformation), we can expand:

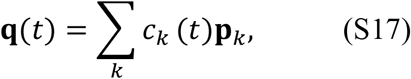

with *c*_*k*_(*t*) = *µ*_*k*_(*t*) in the orthonormal case. In a non-orthogonal basis, the relation involves the Gram matrix, but the essential idea remains: the coefficients *c*_*k*_(*t*) represent the degree to which each engram contributes to the current informational pattern.

#### S2.7. Dynamics with resonant retrieval

The evolution of the coefficients is governed by an equation analogous to (S13) but projected onto the engram basis. Including a nonlinear resonance term yields:

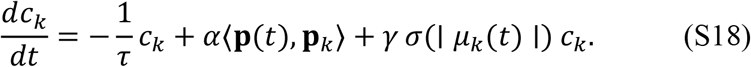

The first term is exponential forgetting, the second is the projection of the sensory input onto the engram, and the third is the resonant amplification that occurs when the overlap exceeds a threshold. The sigmoid *σ* models the threshold nature of memory retrieval: for small overlaps, the term is negligible; for large overlaps, it becomes positive and can overcome forgetting, leading to rapid growth of *c*_*k*_ – i.e., the memory is “brought to mind”.

#### S2.8. Relation to observed microwave power

The microwave emission detected by MWEG is an **epiphenomenon** – a consequence of the fast changes in neural activity that accompany memory retrieval. The power at frequency ω is proportional to the squared magnitude of the time derivative of the relevant coefficients:

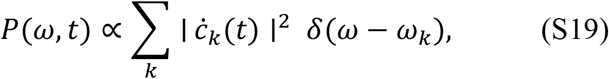

where ω_*k*_ is the characteristic oscillation frequency associated with the *k*-th engram (e.g., the frequency of the neural ensemble that encodes it). This formula shows that the measured spectrum reflects the **dynamics** of informational processes, not the patterns themselves. Thus, by analyzing the spectral peaks, we can infer which memories are being retrieved and how rapidly they change.

#### S2.9. Connection with the energy-based model

The two models are complementary. The energy-based model provides a physical mechanism for microwave generation (coherent oscillations, currents, radiation). The information-based model explains **why** certain modes are excited and why their activity correlates with cognitive states: because the informational overlap *µ*_*k*_(*t*) drives the nonlinear amplification term *γσc*_*k*_, leading to large 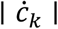 and hence observable spectral peaks. The parameters ω_*k*_ in the energy model correspond to the oscillation frequencies associated with each engram, linking the two descriptions.

### S3. Additional Experimental Results

#### S3.1. Data from the 2021 experiment

The distribution of total substantial bursts during recording from the back of the subject’s head (series 1–5), from the control sample (series 6), and from the background (series 8) of the 2021 study is shown in Figure S3.

**Fig S3.**
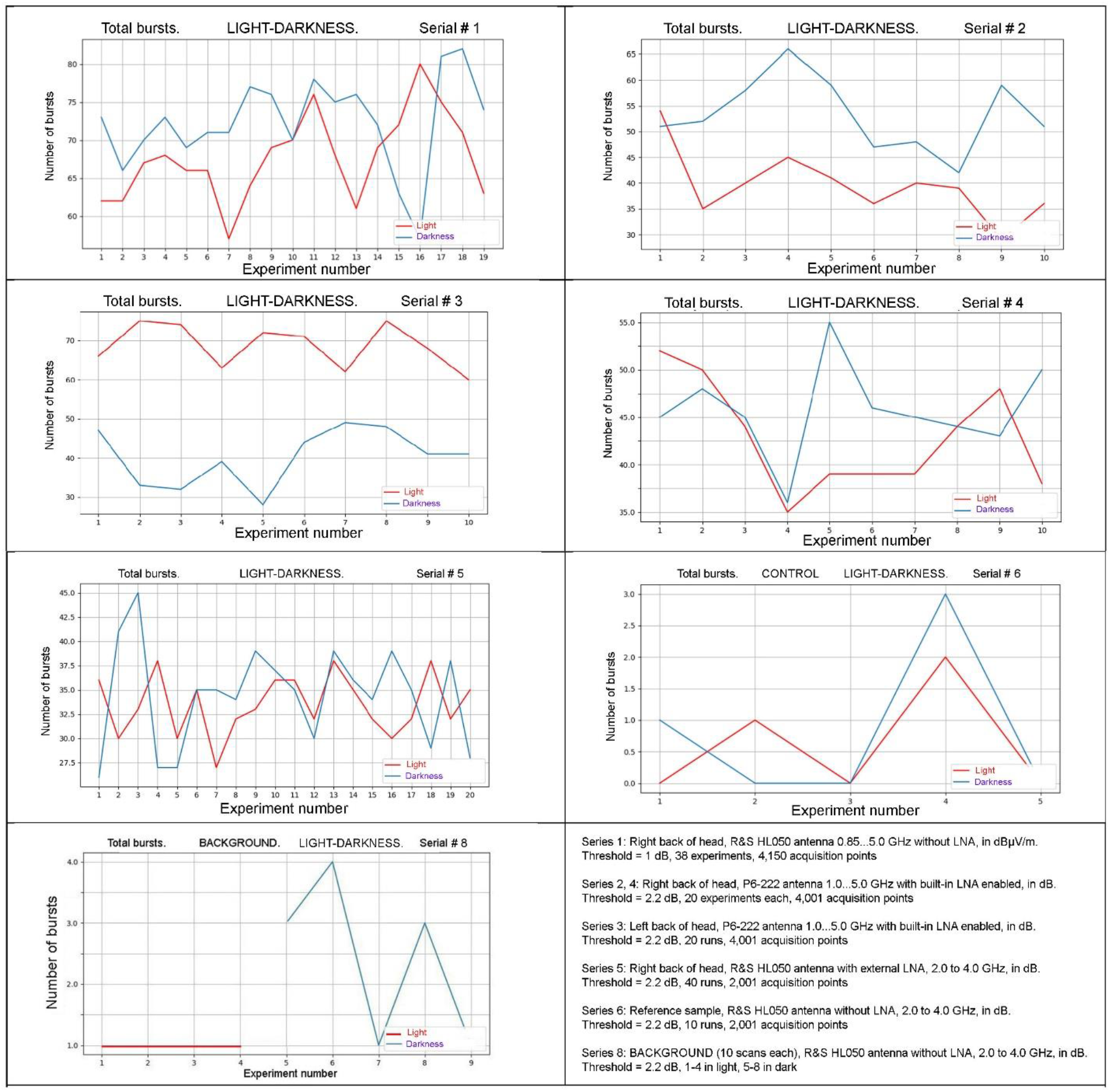
Display of the total significant spikes across all experiments of all series of the 2021 study for the light/dark stimulus.

#### S3.2. Supplementary Figure S4. Frequency distribution of microwave spikes in the 2021 experiment

The histograms on figure S4 show the normalized frequency distribution of all substantial spikes recorded from the right (series 1,2,4,5) and left (series 3) occipital areas under light (red) and darkness (blue) conditions. Data are pooled from 138 measurements from a volunteer, of the 2021 experiment, series 1-5. For the right hemisphere, the distribution shifts towards higher frequencies in darkness compared to light, indicating enhanced high-frequency activity. For the left hemisphere, an opposite trend is observed: higher frequencies occur predominantly under light stimulation. These patterns are consistent with the functional hemispheric asymmetry reported in the main text (Table 2) and support the hypothesis that MWEG captures frequency-specific neural dynamics related to cognitive state.

**Fig S4.**
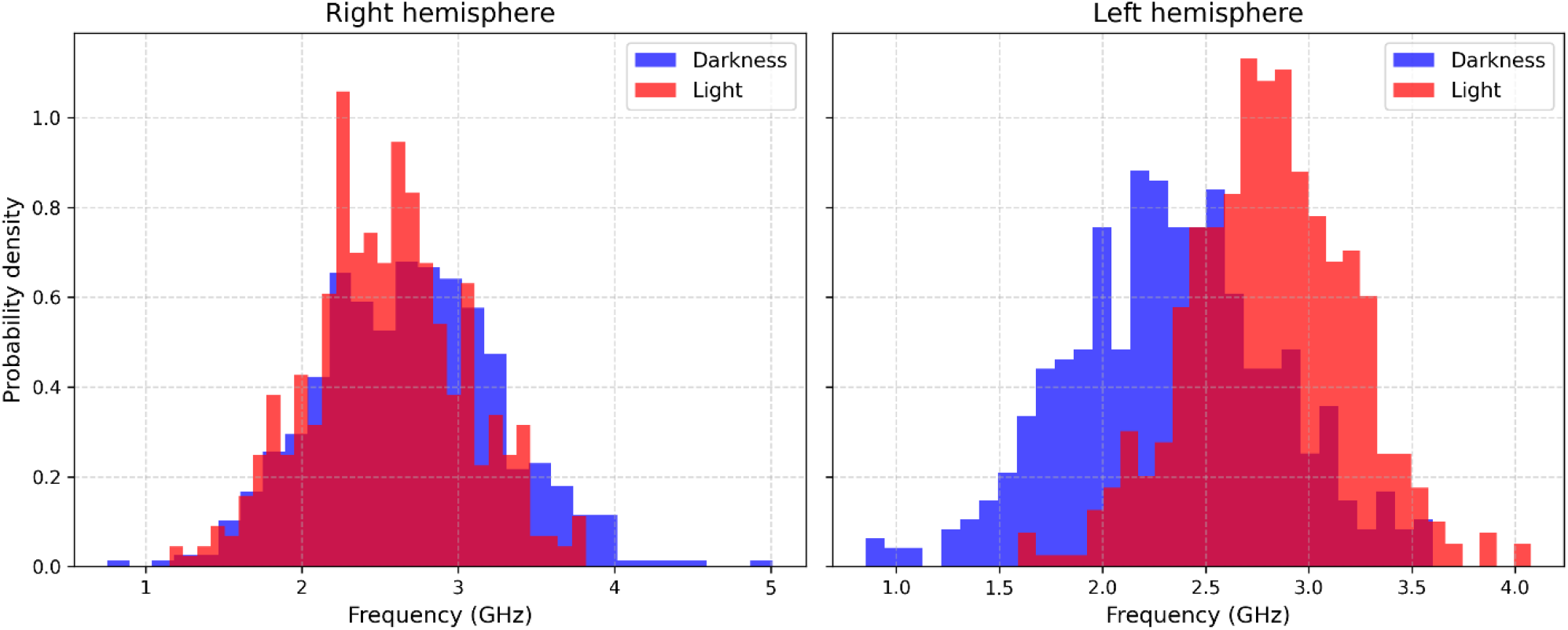
Frequency distribution of microwave spikes in the 2021 experiment: (a) right hemisphere; (b) left hemisphere. Red – light, blue – darkness.

#### S3.3. Supplementary Figure S5. Spike counts across sessions in the 2021 experiment.

The figure 5 shows the number of substantial spikes for each individual measurement in series 1– 5 of the 2021 experiments. Red circles represent light condition, blue squares represent darkness. Each point corresponds to one experimental run. The vertical separation between light and darkness measurements illustrates the reproducibility of the effect across sessions: in series 1, 2 and 3 the darkness values are consistently higher (series 1, 2) or lower (series 3) than the corresponding light values, while series 4 and 5 show overlapping distributions. These visual observations are confirmed by the paired t-test results in Table 2 of the main text. The small circles (fliers) on the boxplots indicate individual outliers that do not affect the overall statistical conclusions

**Fig S5.**
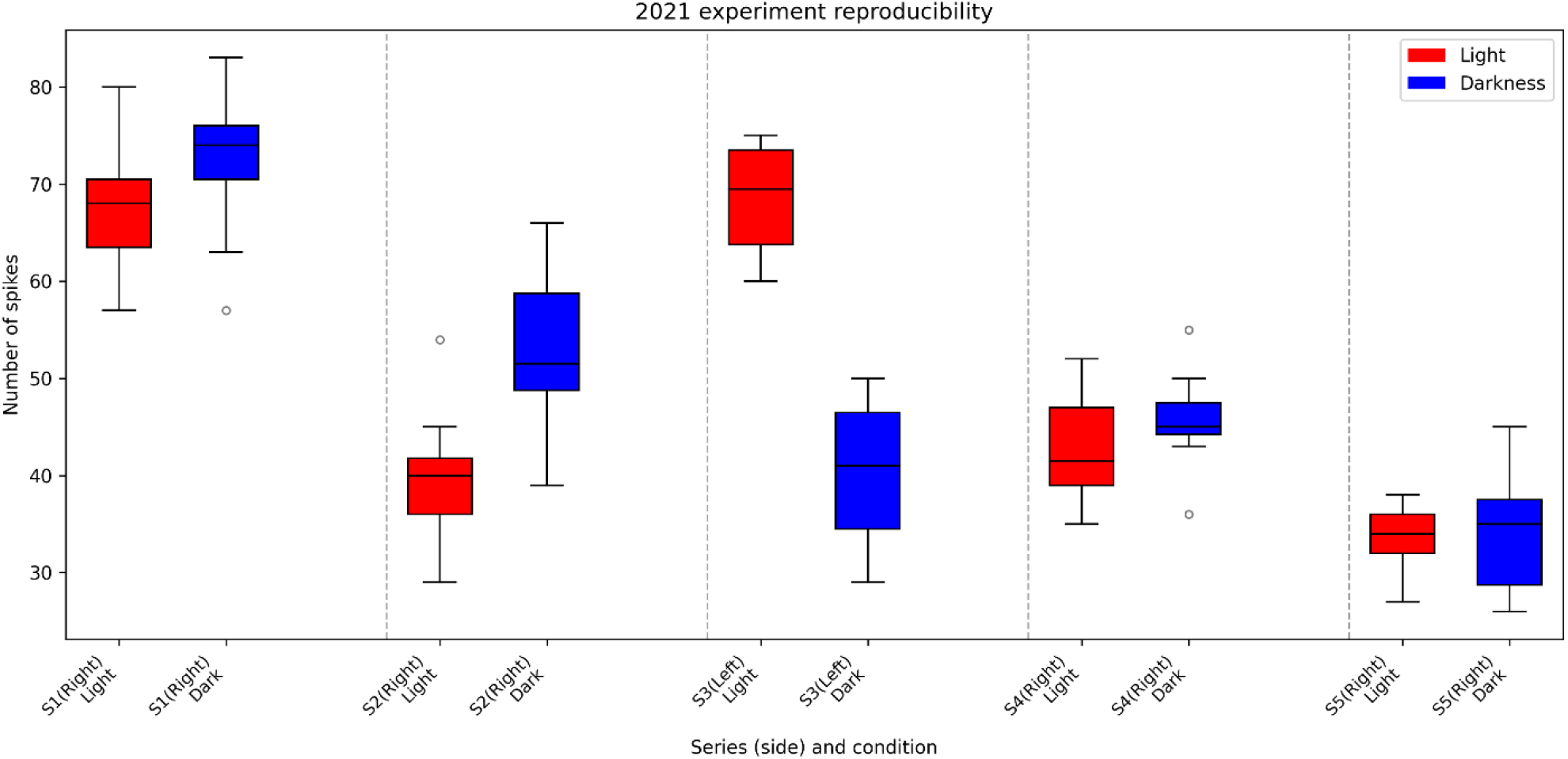
Spike counts across sessions in the 2021 experiment. Each point represents one measurement; circles – light, squares – darkness. The difference is consistent across all sessions.

#### S3.4. Individual subject data from the 2025 experiment

The summary substantial spikes of the 2025 study experiments for the light/dark stimulus, as obtained by the SSPIKES program, are presented in Figure S6.

**Fig S6.**
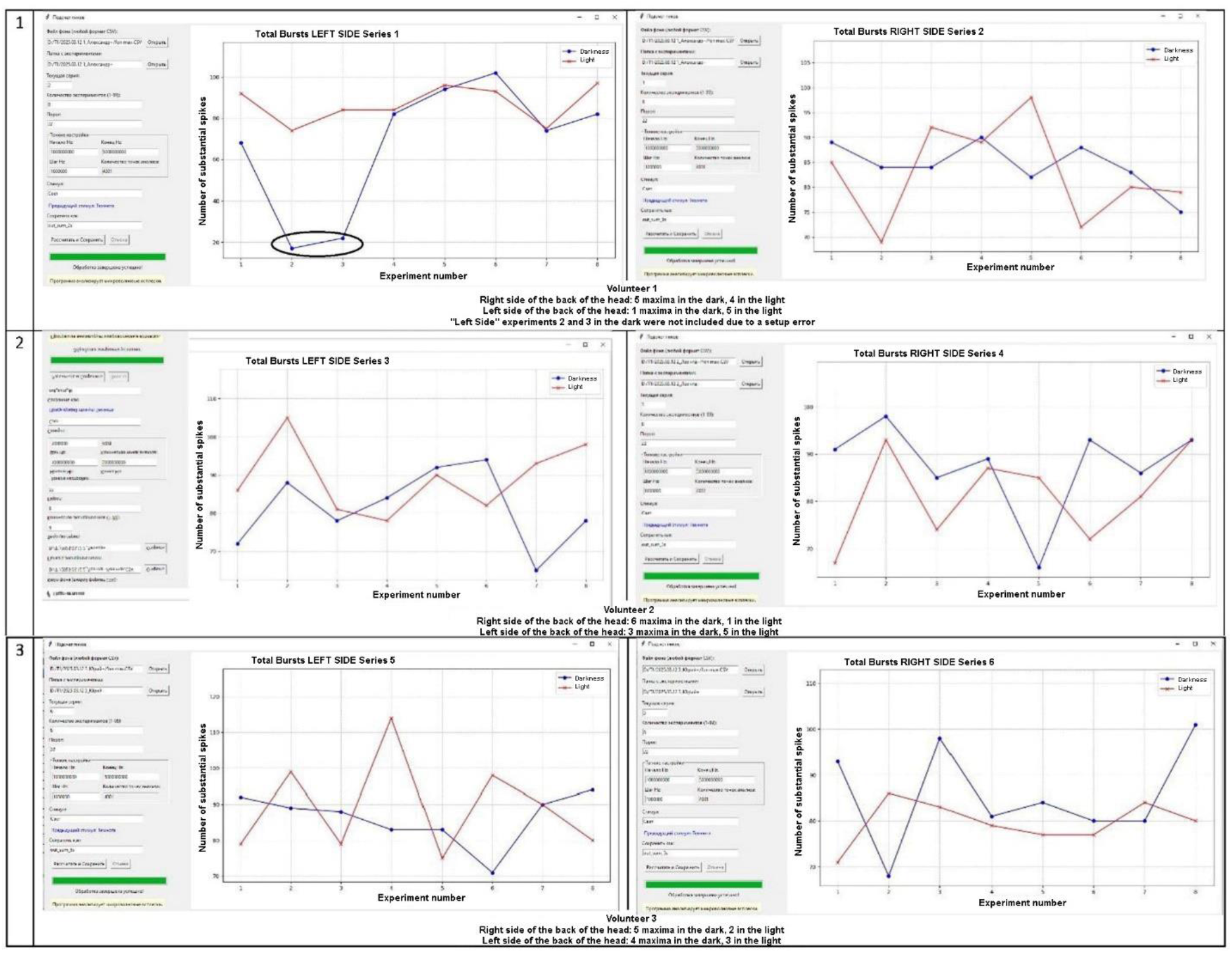
Display of the total significant bursts across all experiments of all series of the 2025 study for the light/dark stimulus.

Table S1 presents the raw spike counts for each of the three subjects in the 2025 experiment, for both hemispheres and both conditions. Data are shown as mean ± standard deviation over the 8 repetitions.

**Table S1.** Individual subject data (2025 experiment)

| Subject | Hemisphere | Light<br>(mean ± SD) | Darkness<br>(mean ± SD) | Difference<br>(D-L) | p<br>(paired t-test) |
| --- | --- | --- | --- | --- | --- |
| S1 | Right | 84.3 ± 4.2 | 86.1 ± 5.1 | +1.8 | 0.12 |
| S1 | Left | 91.2 ± 3.8 | 82.5 ± 4.6 | -8.7 | 0.003 |
| S2 | Right | 81.5 ± 5.0 | 83.8 ± 4.8 | +2.3 | 0.08 |
| S2 | Left | 88.9 ± 4.5 | 80.1 ± 5.2 | -8.8 | 0.004 |
| S3 | Right | 82.9 ± 4.7 | 83.3 ± 5.3 | +0.4 | 0.76 |
| S3 | Left | 88.4 ± 4.1 | 81.6 ± 4.9 | -6.8 | 0.02 |

### S4. Methodological Details

#### S4.1. Calibration and threshold selection

The spike detection threshold was set to 2.2 dB above the averaged background noise floor. This value was chosen empirically based on the distribution of noise in empty-chamber recordings: 2.2 dB corresponds approximately to twice the standard deviation of the noise power, ensuring that fewer than 5% of background fluctuations are falsely counted as spikes. For the 2025 experiment, a higher threshold (22 dB) was used due to the different shielding characteristics of the anechoic shielded chamber, but after calibration the effective sensitivity was comparable; both thresholds correspond to a signal to noise ratio of about 1.6.

#### S4.2. Software implementation

The MWENC program (Python 3.9) reads CSV files exported from the spectrum analyzer, subtracts the averaged background, identifies peaks using scipy.signal.find_peaks with parameters height = threshold, distance = 10 points, and prominence = 1.5 dB. The SSPIKES program aggregates results across experiments and performs statistical tests using scipy.stats.ttest_rel.

#### S4.3. Data availability

All raw data and analysis scripts are available at Zenodo (DOI: 10.5281/zenodo.15739578).

### S5. Discussion of Limitations and Future Directions

#### S5.1. Sample size and generalizability

The 2021 experiment involved a single subject, and the 2025 experiment three. While the effects are statistically robust within this cohort, larger studies are needed to establish normative values and to explore inter-individual variability.

#### S5.2. Spatial resolution

Current recordings use a single antenna, providing no spatial localization. Future work should employ multi-channel antenna arrays (e.g., 8–16 elements) to enable source imaging and to correlate activity with anatomical structures.

#### S5.3. Simultaneous validation with established methods

To definitively link MWEG signals to neural activity, simultaneous recordings with EEG or fMRI are necessary. Such experiments are planned but require careful shielding to avoid interference.

#### S5.4. Theoretical refinements

The mathematical models presented here are first-order approximations. More realistic models would incorporate:

- Nonlinearities in neural dynamics (e.g., Hodgkin–Huxley type equations);
- Detailed geometry of the head for electromagnetic propagation;
- Stochastic fluctuations and noise.

## End of Supplementary Materials

